# Metabolites form a globally connected chemical network across protein families

**DOI:** 10.64898/2026.08.11.744260

**Authors:** Jeffrey Skolnick, Bharath Srinivasan

## Abstract

Metabolites are generally viewed as substrates, products, cofactors, or regulators of individual proteins, whereas metabolites recurring across many protein families are often regarded as promiscuous binders. Here, we analyzed 989,058 BioLiP2 protein–ligand binding sites and assigned 929,546 sites to ECOD v295 homologous groups to quantify ligand specificity, cross-fold scatter, structural breadth, and metabolite-mediated connectivity across protein-family space. Many ancient metabolites preferentially occupied cognate structural groups, demonstrating that broad evolutionary reuse can coexist with local structural discrimination. After excluding elemental metals, BioLiP potential-artifact/dual-use ligands, and metabolites containing fewer than six heavy atoms, 32 ancient metabolites occupied a mean of 185.38 ECOD F-groups per metabolite, compared with 6.32 F-groups for 2,540 mapped filtered non-ancient metabolites— a 29.35-fold enrichment (bootstrap 95% CI, 18.66–43.46). The complete 40-ancient-metabolite network connected all 6,798 associated F-groups into a single giant connected component (GCC). Even after stringent filtering, all 3,135 ancient-metabolite-associated F-groups remained in one GCC. Degree-preserving configuration-model randomizations and maximum-degree capping showed that this connectivity follows from the broad, recurrent distribution of metabolite binding rather than dependence on a few extreme hubs or a specialized higher-order topology. Differences between ancient and filtered non-ancient networks were not explained by metabolite size, whereas generic crystallization additives preferentially occupied smaller pockets. These results indicate that a limited ancient chemical repertoire established a globally connected protein-family architecture that subsequent metabolite diversification expanded while preserving its basic organization.

**Significance:** Metabolites are conventionally viewed as substrates, products, cofactors, or regulators acting on individual proteins. Global examination of experimentally observed metabolite–protein interactions reveals a broader organizing principle. Ancient metabolites combine local binding discrimination with extraordinary reuse across protein families, such that only 40 metabolites generate an almost completely connected network spanning thousands of ECOD (evolutionary classification of domains) protein families. The much larger non-ancient metabolite repertoire expands the protein-family space covered by this network, while preserving near-global connectivity. Thus, metabolite diversification appears to have elaborated, rather than created, a chemically connected protein architecture established early in evolution, suggesting that overlapping metabolite-binding repertoires could coordinate proteins, pathways, and cellular processes.

## Introduction

Proteome-wide interaction maps show that metabolites influence enzyme activity, allostery, protein-complex remodeling, localization, phase behavior, protein stability, signaling, and gene expression beyond their canonical roles as substrates and products (1, 2). If a limited set of central metabolites reproducibly binds defined subsets of proteins, overlapping binding repertoires could provide a chemical means of coupling protein and pathway behavior (3). Cross proteome metabolite–protein binding is the physical mechanism underlying the Entabolon hypothesis, in which shared metabolite binding is conjectured to correlate protein behavior and give rise to protein pathways (3). The central objections to this idea are: 1. apparent promiscuity: ancient cofactors and central metabolic intermediates are among the most highly reused ligands in the structural record and are therefore often assumed to bind indiscriminately. 2. binding does not necessarily translate into biochemical function. Nevertheless, binding is the minimal requirement for such functional modification to occur. Here, we ask how globally connected the space of all proteins is due to their possible binding of common metabolites. If the resulting giant component of the metabolite protein -interaction network covers almost all protein families, this would establish the minimal requirement of the Entabolon hypothesis and set the stage for subsequent experimental validation of the functional consequences of such global network correlation.

Two distinctions are essential. First, ligands that are found in many crystallized proteins are mechanistically heterogeneous. Glycerol, polyethylene glycol, and 2-methyl-2,4-pentanediol are largely nonspecific, whereas imidazole and citrate favor broad classes of metal- or anion-binding sites; sulfate, phosphate, and related oxyanions can be both crystallization components and biologically meaningful ligands (see Table S1). Second, broad metabolite reuse need not imply random occupancy of structural space. A metabolite can bind many proteins while remaining concentrated in a cognate ECOD superfamily or a restricted set of structural families (4). Because specific interactions have finite information capacity (5), the relevant question is not simply how many proteins a metabolite binds, but how nonuniformly those interactions are distributed across protein structural space. However, if an endogenous metabolite binds many proteins (implying a favorable binding free energy), it cannot simply be dismissed as a crystallization artifact. We therefore performed a whole-PDB census (6) to ask whether ancient metabolites (7) preferentially occupy cognate structural groups, whether their apparent cross-fold scatter is explained by ligand size, and whether their overlapping binding repertoires generate higher-order connectivity across protein space. We then extended this analysis to non-ancient metabolites as well to see what differentiates the binding pattern of ancient from non-ancient metabolites.

## Results

### Ligand size explains much apparent cross-fold promiscuity

Cross-fold scatter decreases monotonically with heavy-atom count across all ligands (Spearman ρ (8) = −0.29, P = 3.8 × 10⁻²⁵). Class-level size statistics are summarized in Table 1.

**Table 1.**
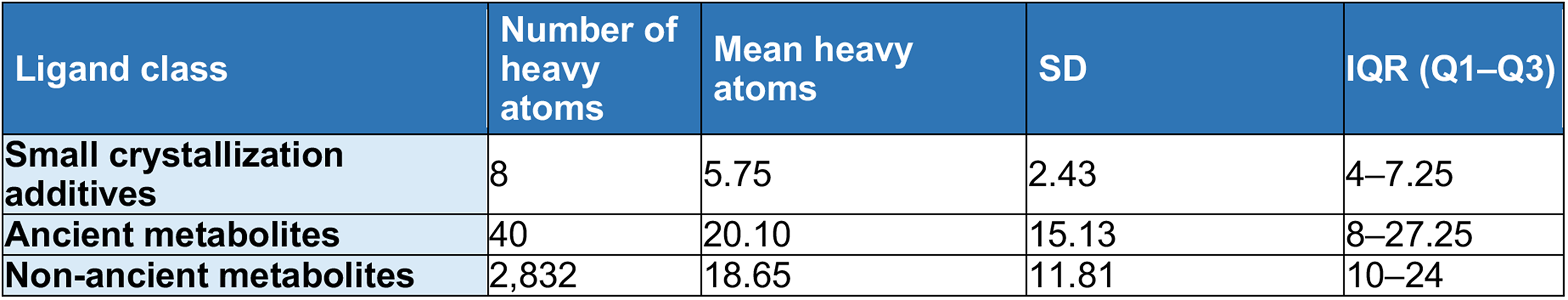
Summary of ligand molecular size measured by number of non-hydrogen (heavy) atoms.

| Ligand class | Number of heavy atoms | Mean heavy atoms | SD | IQR (Q1–Q3) |
| --- | --- | --- | --- | --- |
| Small crystallization additives | 8 | 5.75 | 2.43 | 4–7.25 |
| Ancient metabolites | 40 | 20.10 | 15.13 | 8–27.25 |
| Non-ancient metabolites | 2,832 | 18.65 | 11.81 | 10–24 |

The small crystallization additives contain 5.75 ± 2.43 heavy atoms (IQR, 4–7.25), compared with 20.10 ± 15.13 (IQR, 8–27.25) for the 40 ancient metabolites and 18.65 ± 11.81 (IQR, 10–24) for 2,832 non-ancient metabolites. Such molecules present fewer steric and chemical constraints and can therefore more readily bind to a diverse set of small protein pockets. We also tested an initial geometric explanation based on pocket depth, but the apparent difference between specific and broadly binding ligands disappeared after controlling for unequal site representation. The size–scatter relationship persisted, whereas pocket depth did not explain cross-fold scatter. Thus, much apparent ligand promiscuity is an expected structural consequence of small molecular size rather than an intrinsic property of metabolites.

### Pocket-volume analysis and ligand-balanced rarefaction

Pocket-volume distributions were grouped into specific metabolites, generic crystallization additives (7) and the broader ancient-metabolite set (see Tables S1 and S2). Because the number of PDB observations differed substantially among ligands, class-level comparisons based on all pockets could be dominated by highly represented ligands. We therefore performed ligand- balanced rarefaction at sampling depths of 5, 10, 15, and 20 pocket observations per ligand. At each depth, ligands with at least the required number of observations were retained; the specified number of pocket volumes was sampled independently from each ligand without replacement and pooled within ligand class. The class median pocket volume was calculated, and the entire procedure was repeated 2,000 times using fixed pseudorandom seeds for reproducibility. Class estimates are reported as the median of the 2,000 replicate class medians, with uncertainty summarized by the 2.5th and 97.5th percentiles of the replicate distribution. Pairwise class differences were evaluated within each replicate using two-sided Mann–Whitney U tests (9), and U/(n₁n₂) was used as a probability-of-superiority effect size. As an independent analysis that treated the ligand rather than the individual pocket as the unit of observation, the median pocket volume was calculated separately for each ligand and distributions of ligand-specific medians were compared between classes using two-sided Mann–Whitney U tests (10). Concordance across rarefaction depths and with the ligand-level analysis was used to assess robustness to unequal ligand representation and pocket-level pseudo-replication.

Pocket volume did not explain the distinction between fold-specific and broadly binding ancient metabolites. Instead, it sharply separated metabolite-containing binding sites from those occupied by generic crystallization additives (see Table 2). To control for the highly unequal number of PDB observations per ligand, we performed ligand-balanced rarefaction at depths of 5, 10, 15, and 20 examples of pockets per ligand. Across all four sampling depths, the median rarefied pocket volume remained near 2,000 Å³ for both specific metabolites and the broader ancient-metabolite/additive set, whereas generic additives occupied substantially smaller pockets (∼665–673 Å³). At 20 pockets per ligand, median rarefied pocket volumes were 2,058 Å³ for specific metabolites, 2,090 Å³ for the ancient-metabolite/additive set, and 665 Å³ for generic additives. The same conclusion was obtained when the ligand, rather than the individual pocket, was treated as the unit of observation: median ligand-specific pocket volumes were 2,068, 2,280, and 774 Å³, respectively. Specific metabolites and ancient metabolites/additives each differed from generic additives (two-sided Mann–Whitney U (9), P = 0.0040 and P = 0.0043, respectively), whereas the two metabolite-containing classes did not differ (P = 0.691). Thus, the pocket-volume separation is not driven by highly represented ligands or pocket-level pseudo-replication. Generic additives preferentially sample smaller, permissive cavities, whereas ancient metabolites more often occupy larger, structurally elaborated pockets compatible with selective molecular recognition.

**Table 2.**
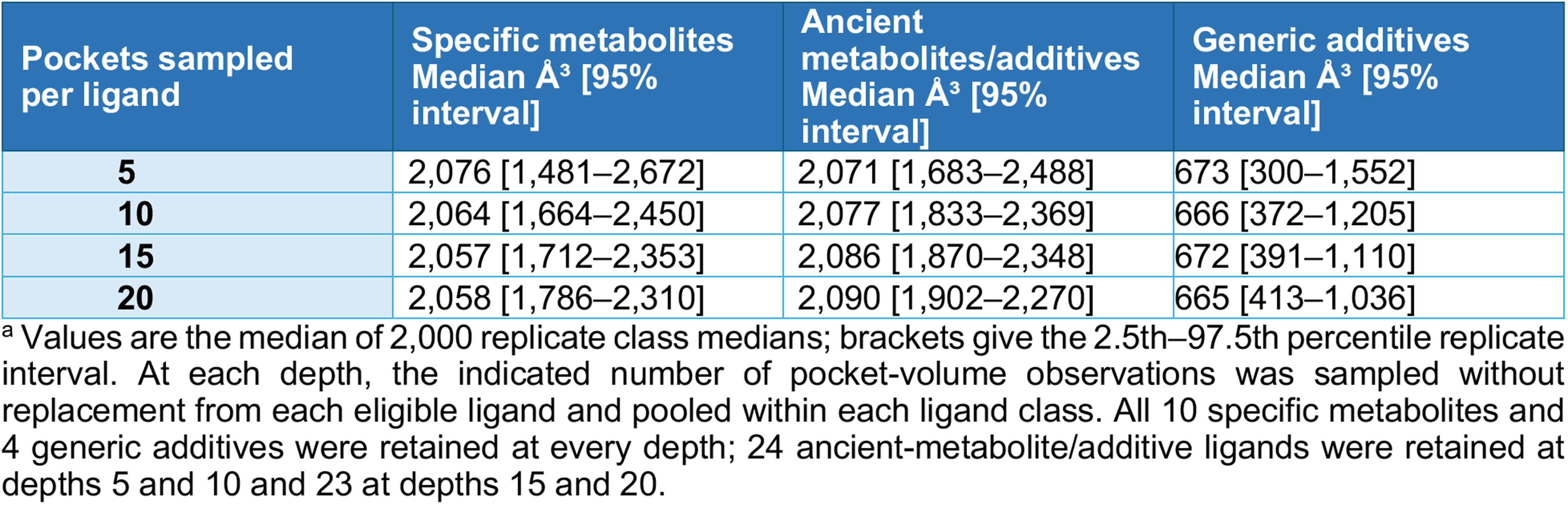
Pocket-volume separation is robust to ligand-balanced rarefaction^a^.

| Pockets sampled per ligand | Specific metabolites<br>Median Å<sup>3</sup> [95% interval] | Ancient metabolites/additives<br>Median Å<sup>3</sup> [95% interval] | Generic additives<br>Median Å<sup>3</sup> [95% interval] |
| --- | --- | --- | --- |
| 5 | 2,076 [1,481–2,672] | 2,071 [1,683–2,488] | 673 [300–1,552] |
| 10 | 2,064 [1,664–2,450] | 2,077 [1,833–2,369] | 666 [372–1,205] |
| 15 | 2,057 [1,712–2,353] | 2,086 [1,870–2,348] | 672 [391–1,110] |
| 20 | 2,058 [1,786–2,310] | 2,090 [1,902–2,270] | 665 [413–1,036] |

<sup>a</sup> Values are the median of 2,000 replicate class medians; brackets give the 2.5th–97.5th percentile replicate interval. At each depth, the indicated number of pocket-volume observations was sampled without replacement from each eligible ligand and pooled within each ligand class. All 10 specific metabolites and 4 generic additives were retained at every depth; 24 ancient-metabolite/additive ligands were retained at depths 5 and 10 and 23 at depths 15 and 20.

### Whole-PDB census of metabolite–fold occupancy

We analyzed the full 2026 BioLiP2 annotation set, comprising 989,058 binding sites (12, 13), and joined it to ECOD v295 domains (5) by residue-range attribution. Of these sites, 929,546 (94.0%) were assigned to an ECOD H-group. ECOD H-groups, which represent homologous superfamilies, were used to quantify site-level specificity and cross-fold scatter. ECOD F-groups, the more specific family level, were used independently to quantify structural breadth and construct the fold-family networks. Among the ECOD v295 F-groups represented in BioLiP2, 9,833 had qualifying experimentally observed metabolite-binding associations. Absence from this set does not imply that other F-groups cannot bind metabolites; it indicates only that they lacked a qualifying association in the datasets analyzed here. For each ligand, specificity was the fraction of assigned sites in its modal H-group, cross-fold scatter was the effective number of occupied H-groups [exp(Shannon entropy)] rarefied to a fixed site count, and molecular size was the number of nonhydrogen atoms in the PDB Chemical Component Dictionary. The 40 ancient metabolites represented in the final HMDB/ChEBI–BioLiP–ECOD intersection are listed in Table S3.

### Ancient metabolites mostly concentrate in their cognate folds

Many ancient metabolites have a clear home fold (Table 3). Thiamine diphosphate is nearly fold-restricted, with 98% of assigned sites in its modal fold. NAD and 2-oxoglutarate are also strongly concentrated, with specificities of 0.76 and 0.64, respectively. The nucleotide-binding Rossmann fold (11) provides the canonical example; it is the dominant structural context for NAD and for SAM/SAH-dependent methyltransferases.

**Table 3.** Cognate-fold specificity of fold-specific ancient metabolites^a^.

| Metabolite | Cognate (modal) ECOD fold | Specificity | Heavy atoms |
| --- | --- | --- | --- |
| **NAD** | Rossmann (NAD-binding) | 0.76 | 44 |
| **Thiamine diphosphate** | thiamine-diphosphate fold | 0.98 | 26 |
| **2-oxoglutarate** | DSBH dioxygenase (Fe/2OG) | 0.64 | 10 |
| **SAM / SAH** | Rossmann (class I methyltransferase) | 0.49–0.77 | 26–27 |
| **GDP** | Rossmann-related (sugar-nucleotide) | 0.62 | 28 |
| **PLP** | PLP-dependent transferase | 0.64 | 16 |

<sup>a</sup> Specificity is the fraction of assigned binding sites in the modal ECOD H-group, calculated from the BioLiP2–ECOD v295 join of 929,546 assigned sites. Values differ modestly from earlier estimates for 2-oxoglutarate (0.64 versus 0.76) and PLP (0.64 versus 0.53) because of ECOD H-group boundaries. GDP is concentrated in the sugar-nucleotide Rossmann set rather than the P-loop, which accounts for a minority of GDP sites in this PDB census. Complete per-metabolite data are provided in Table S2.

Not all ancient metabolites are equally restricted. Nucleotides, phosphate, and metals occupy a broader range of folds, but much of this breadth reflects repeated binding within broad, defined fold families rather than random cross-fold occupancy (Table 4). ATP and ADP recur predominantly within P-loop NTPases and kinases (specificities 0.54 and 0.49), whereas GTP is concentrated in Rossmann-related and GTPase folds (0.65). Inorganic phosphate repeatedly occupies phosphate- and anion-binding pockets, and catalytic metals recur in metal-dependent enzymes; the latter constitute the local-modulator class examined in our companion paper. Thus, at the superfamily level, many apparently broad-binding metabolites are better described as broad-family anchors than as generalists. This is the same distinction that separates true crystallization artifacts from fold-biased additives (see Introduction and Table S1). Only the smallest ligands lack a preferred structural family, and these are predominantly additives rather than metabolites.

**Table 4.** Broadly binding ancient metabolites and their cross-fold occupancy^a^.

| Metabolite | Favored broad fold family | Specificity | Scatter | Heavy atoms |
| --- | --- | --- | --- | --- |
| **ATP** | P-loop NTPase / kinases | 0.54 | 6.9 | 31 |
| **ADP** | P-loop NTPase / kinases | 0.49 | 5.9 | 27 |
| **AMP** | mononucleotide-binding | 0.10 | 21.3 | 23 |
| **GTP** | Rossmann / GTPase (mixed) | 0.65 | 3.6 | 32 |
| **CoA / acetyl-CoA** | multiple (broad) | 0.18 / 0.41 | 12.2 / 7.5 | 48 / 51 |
| **FAD** | FAD-binding oxidoreductases (Rossmann) | 0.50 | 5.7 | 53 |
| **NADP** | Rossmann (dehydrogenases) | 0.63 | 3.5 | 48 |
| **Mg<sup>2+</sup>** | metal-dependent enzymes (very broad) | 0.09 | 25.8 | 1 |
| **Zn<sup>2+</sup>** | metalloenzymes / Zn-fingers | 0.10 | 24.5 | 1 |

<sup>a</sup> Scatter is the effective number of occupied H-groups [ $\exp(\text{Shannon entropy})$ ], rarefied to 50 sites over 30 replicates. ATP, ADP, GTP, NADP, and FAD remain broad-family anchors (specificity, 0.49–0.65; scatter, 3.5–6.9), whereas AMP, CoA, and metals show substantially broader occupancy (specificity, 0.09–0.18; scatter, 12–26). For the anchor class, apparent breadth largely reflects repeated binding within a broad, defined superfamily—for example, ATP binding to P-loop NTPases rather than random cross-fold scatter. Metals represent the size extreme of one heavy atom.

### Ancient-metabolite binding connects protein families

Projection of ancient-metabolite binding onto ECOD F-groups further revealed a densely interconnected architecture (Figure 1). Nodes represent ECOD F-groups and are scaled by the number of distinct ancient metabolites represented; edges connect families sharing ancient metabolites. The network shows how a limited chemical vocabulary can be reused across diverse structural families without requiring uniform, indiscriminate binding. Local H-group preference and global F-group connectivity are complementary properties: selective binding at the molecular scale, combined with exceptional metabolite reuse, is sufficient to generate system-wide connectivity.

**Figure 1.**
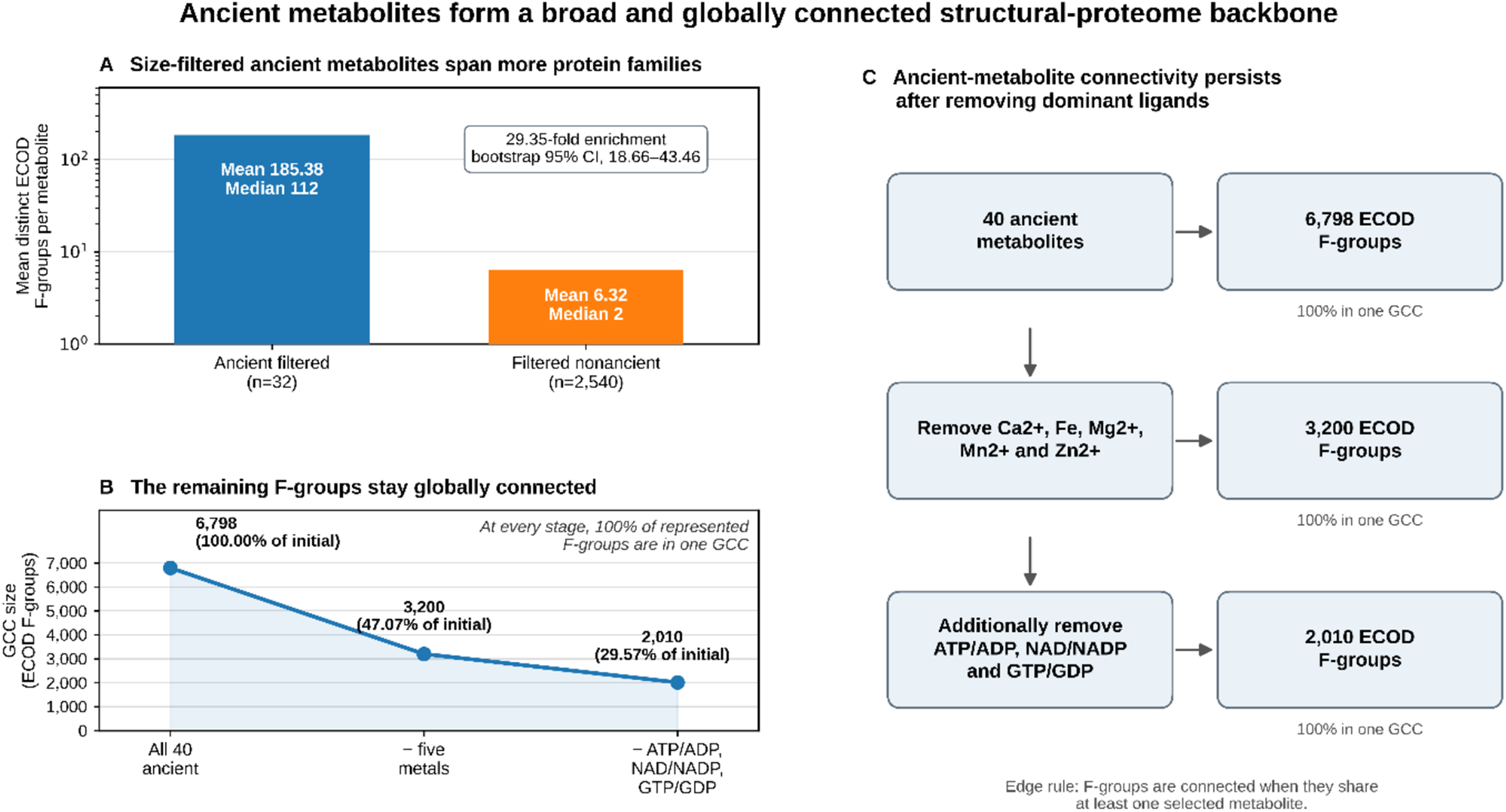
Ancient metabolites span and connect protein structural space. (A) Mean numbers of distinct ECOD F-groups associated with 32 stringently filtered ancient metabolites and 2,540 mapped stringently filtered non-ancient metabolites; medians are annotated. Ancient metabolites exhibit 29.35-fold greater mean F-group breadth (bootstrap 95% CI, 18.66–43.46). (B) The complete 40-ancient-metabolite network contains 6,798 ECOD F-groups. After removal of the five represented elemental metals, 3,200 F-groups remain; after additional removal of ATP/ADP, NAD/NADP, and GTP/GDP, 2,010 remain. At each stage, 100% of the represented F-groups belong to one GCC. (C) Schematic of the same sequential-removal analysis. F-groups are connected when they share at least one retained ancient metabolite.

### Ancient metabolites define a giant component spanning protein families

We next asked whether this structural breadth produces isolated modules or a globally connected protein network. We reconstructed the protein–metabolite graph de novo from the BioLiP2 annotation file using the 40 ancient metabolites represented in the fold-breadth analysis. Protein nodes were PDB chains and were connected in the one-mode projection when they shared at least one ancient metabolite. The resulting graph contained 189,201 protein chains, all of which belonged to a single giant connected component (GCC). Removal of the five represented metal ions (Ca²⁺, Fe, Mg²⁺, Mn²⁺, and Zn²⁺) left 83,270 chains. Additional exclusion of ATP/ADP, NAD/NADP, and GTP/GDP left 47,112 chains. At each removal stage, all retained chains belonged to one GCC. Thus, complete chain-level connectivity persisted after sequential removal of the dominant metals and nucleotides.

Very small ancient metabolites and metal ions accounted for much of the breadth of protein-family coverage, but this expansion was dominated by biologically functional metal ions rather than small organic metabolites. Of the 6,798 ECOD F-groups associated with ancient metabolites, 3,661 (53.9%) were observed exclusively with ancient ligands containing five or fewer heavy atoms. These families were predominantly associated with Zn²⁺ (1,794 families), Ca²⁺ (1,518), and Mg²⁺ (1,337), whereas glycine was associated with only 63 families. Alanine was excluded from this category because its corrected heavy-atom count is six. Zn²⁺, Ca²⁺, and Mg²⁺ uniquely accounted for 1,084, 779, and 523 families, respectively. Overall, 69.3% of these small-ligand-associated families bound only one of the six ≤5-heavy-atom ancient species, whereas 30.7% bound two or more. Because these interactions were obtained from the biologically curated BioLiP2 dataset, their contribution cannot simply be attributed to nonspecific occupancy. Removal of the five elemental metals reduced the active family network from 6,798 to 3,200 F-groups, all of which remained in one GCC. Applying the complete stringent filter, which also removed BioLiP potential-artifact/dual-use ligands and all compounds containing fewer than six heavy atoms, left 32 ancient metabolites connecting 3,135 F-groups, again in a single GCC. Thus, small functional ligands expand the reach of ancient chemical space, whereas chemically larger ancient metabolites independently preserve global protein-family connectivity.

Figure 2 shows the strikingly high node-degree distribution of the ancient-metabolite network. The key question is whether the enormous degree of the ≥1 shared ancient-metabolite network is simply caused by closely related proteins being split among different ECOD F-groups. To address this issue, we calculated the mean and median maximum sequence identity for edges sharing 1, 2, 3, 4, 5, and ≥ 6 ancient metabolites, with confidence intervals. Even using an upward-biased “maximum identity” estimate, the median maximum sequence identity is only 33.9–36.4% across edge-weight bins. Edges with >50% identity are approximately 0.006% of the entire set and are therefore negligible. We further discuss this point below.

**Figure 2.**
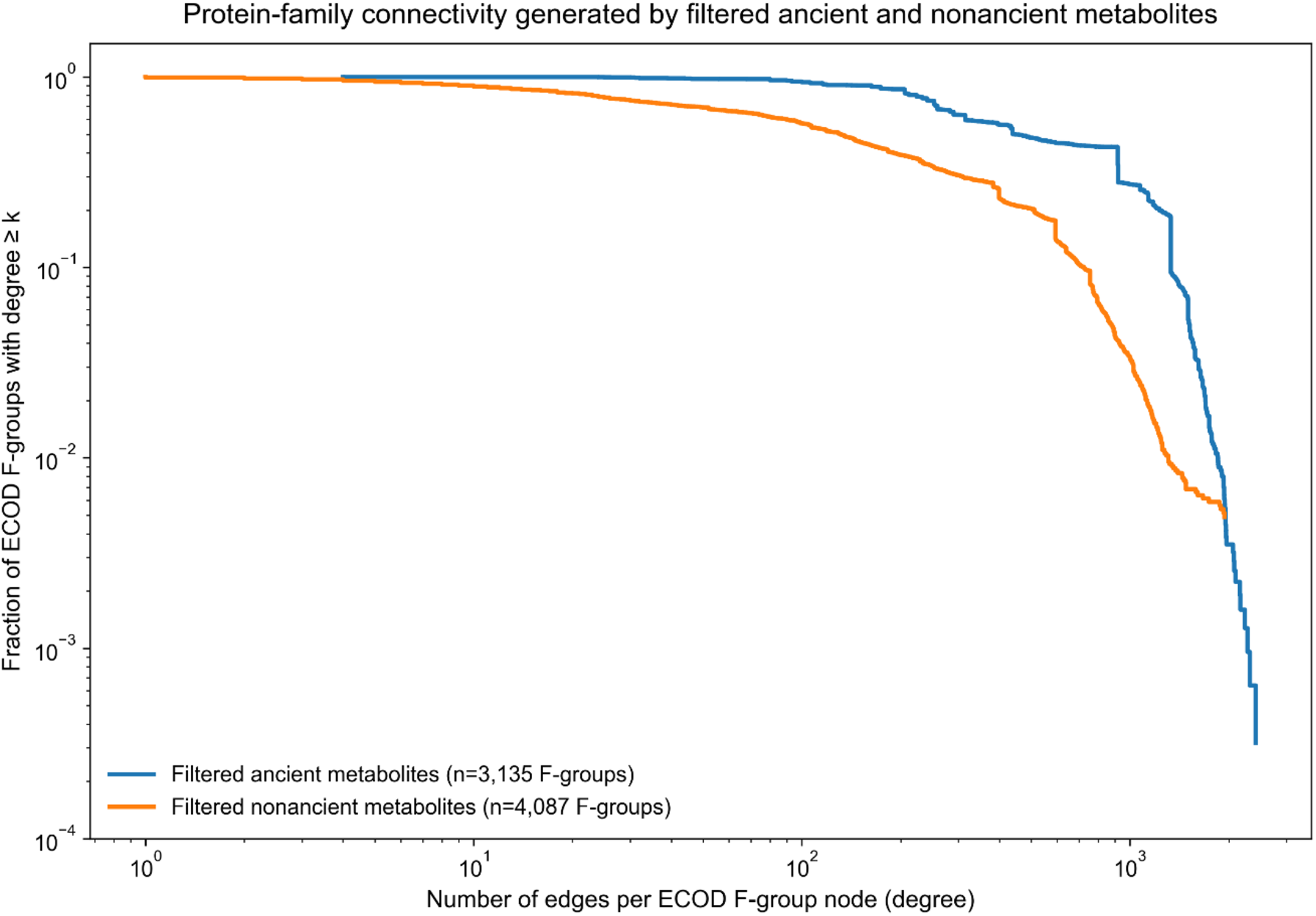
Protein-family connectivity generated by ancient and filtered non-ancient metabolites. Complementary cumulative degree distributions show the fraction of ECOD F-groups with degree ≥ k, where degree is the number of other F-groups connected to a node by at least one shared metabolite.

We first tested higher-order organization at the protein-chain level. The protein–metabolite bipartite graph was randomized by degree-preserving edge swaps (15), thereby retaining both the number of ancient metabolites bound by each chain and the number of chains associated with each metabolite. Across 412 randomized networks, the observed GCC fractions fell within the empirical null distributions at every removal step. For the complete network, the null mean GCC fraction was 99.9995% (95% empirical interval, 99.9954–100%), compared with 100% observed. After metal removal, the null mean was 99.9916% versus 99.9897% observed; after additional ATP/ADP removal, 99.9843% versus 99.9868%; after NAD/NADP removal, 99.9776% versus 99.9847%; and after GTP/GDP removal, 99.9551% versus 99.9790%. None differed significantly from the chain-level degree-preserving null (all empirical two-sided P ≥ 0.57).

We separately evaluated the ECOD F-group network using the observed family node-degree distribution, progressively capped the maximum degree permitted for each node, and reconstructed 50 randomized configuration-model networks at each degree cap (Figure 3). The observed network contained 6,798 nodes and 10,011,118 edges, with a mean degree of 2,945.3, a median degree of 2,871, and a GCC containing 100% of the nodes. In the family-level null, the GCC remained 6,798/6,798—100%—in every randomization when maximum degree was capped at five or three edges per node. When every node was capped at degree two, the mean GCC decreased to 4,913 nodes (72.3%), with substantial replicate-to-replicate variability (SD, 1,417 nodes; range, 2,106–6,792) because the randomized degree-two networks decomposed into cycles. At degree one, the GCC collapsed to pairs. Thus, neither the chain-level nor the family-level GCC required a special higher-order topology or the extraordinary hubs of the observed network; in the capped family-level null, three connections per family were sufficient to connect all 6,798 families.

**Figure 3.**
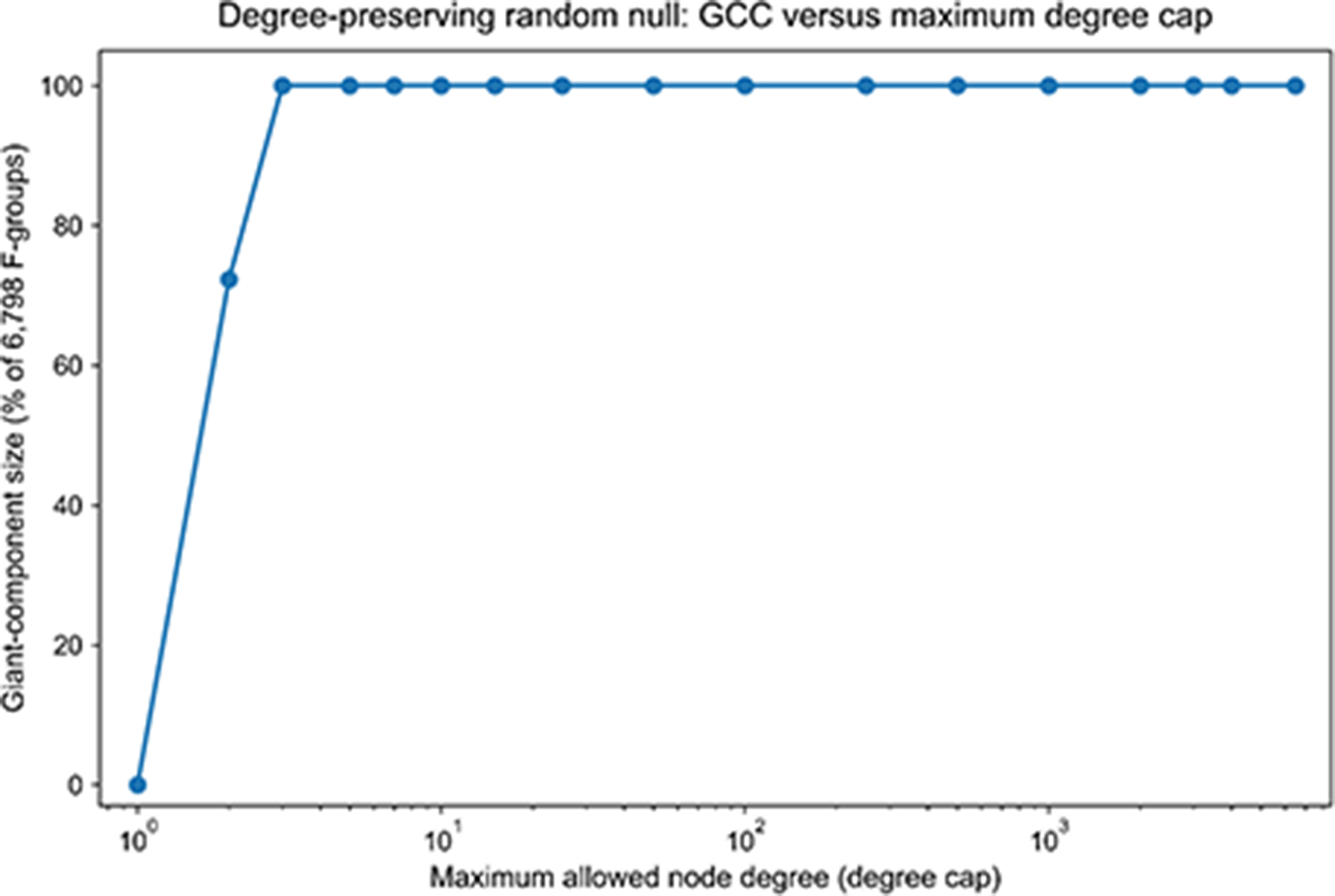
Size of the GCC versus the maximum allowed degree per node in the ancient-metabolite network.

As noted above, when an edge was defined by at least one shared ancient metabolite, the network contained 6,798 ECOD F-groups, all of which belonged to a single giant connected component. The network contained 10,011,118 unique edges, with a mean node degree of 2,945.3 and a median degree of 2,871. Tightening the edge criterion to require at least two distinct shared metabolites reduced the active network to 3,603 F-groups, representing 53.00% of all ancient-metabolite-associated F-groups, and 1,822,740 qualifying edges. Nevertheless, all 3,603 active F-groups remained within a single connected component. Thus, the 40 ancient metabolites alone were sufficient to establish a globally connected network spanning a broad and structurally diverse collection of ECOD F-group protein families.

### Network connectivity of non-ancient metabolites

Removal of the BioLiP potential-artifact/dual-use ligand class, followed by exclusion of elemental metals and compounds containing fewer than six heavy atoms, markedly reduced the apparent breadth and density of the non-ancient metabolite–protein-family network. Of 2,892 CCD-resolved non-ancient metabolites, 178 overlapped the BioLiP exclusion list, leaving 2,714 metabolites. After applying the heavy-atom filter and excluding compounds with unresolved sizes, 2,556 non-ancient metabolites remained, of which 2,540 mapped to ECOD F-groups. The resulting network contained 4,087 ECOD F-groups. When an edge required at least one shared metabolite, 4,055 families (99.22%) belonged to the GCC. Requiring at least two shared types of metabolites reduced the GCC to 2,154 families, corresponding to 52.70% of all non-ancient- metabolite-associated families and 95.31% of the 2,260 families retaining at least one qualifying edge. Thus, even after stringent filtering, metabolites in general generated extensive protein-family connectivity, although the connectivity produced by the compact ancient repertoire was substantially denser.

The filtered non-ancient network had a mean node degree of 251.6 and a median degree of 130, compared with 678.6 and 442, respectively, for the network generated by the 32 filtered ancient metabolites. The ancient set therefore generated 2.70-fold greater mean node connectivity despite containing approximately 80-fold fewer mapped metabolites. Although the ancient network contained fewer total F-groups than the non-ancient network—3,135 versus 4,087—it reached 76.7% of the non-ancient family coverage with only 1.26% as many metabolites. On a per-metabolite basis, the mean F-group breadth was 185.4 for ancient metabolites and 6.32 for filtered non-ancient metabolites, representing a 29.35-fold enrichment for the ancient set.

In Figure 4, metabolites are grouped by the number of heavy atoms, and bar heights show the percentage of each class within a size bin. After exclusion of BioLiP potential-artifact/dual-use ligands, elemental metals, and compounds containing fewer than six heavy atoms, the analysis included 32 ancient and 2,556 non-ancient metabolites. Ancient metabolites contained 23.25 ± 13.35 heavy atoms (median, 23; IQR, 10.75–31), compared with 19.53 ± 11.23 heavy atoms (median, 17; IQR, 11–25) for filtered non-ancient metabolites. The distributions did not differ significantly by Welch’s *t* test (*t* = 1.571, df = 31.55, *P* = 0.126), Mann–Whitney *U* test (*U* = 47,101.5, *P* = 0.139), or two-sample Kolmogorov–Smirnov test (*D* = 0.214, *P* = 0.095). Thus, the greater per-metabolite protein-family breadth of ancient metabolites cannot be explained by a gross difference in molecular size.

**Figure 4.**
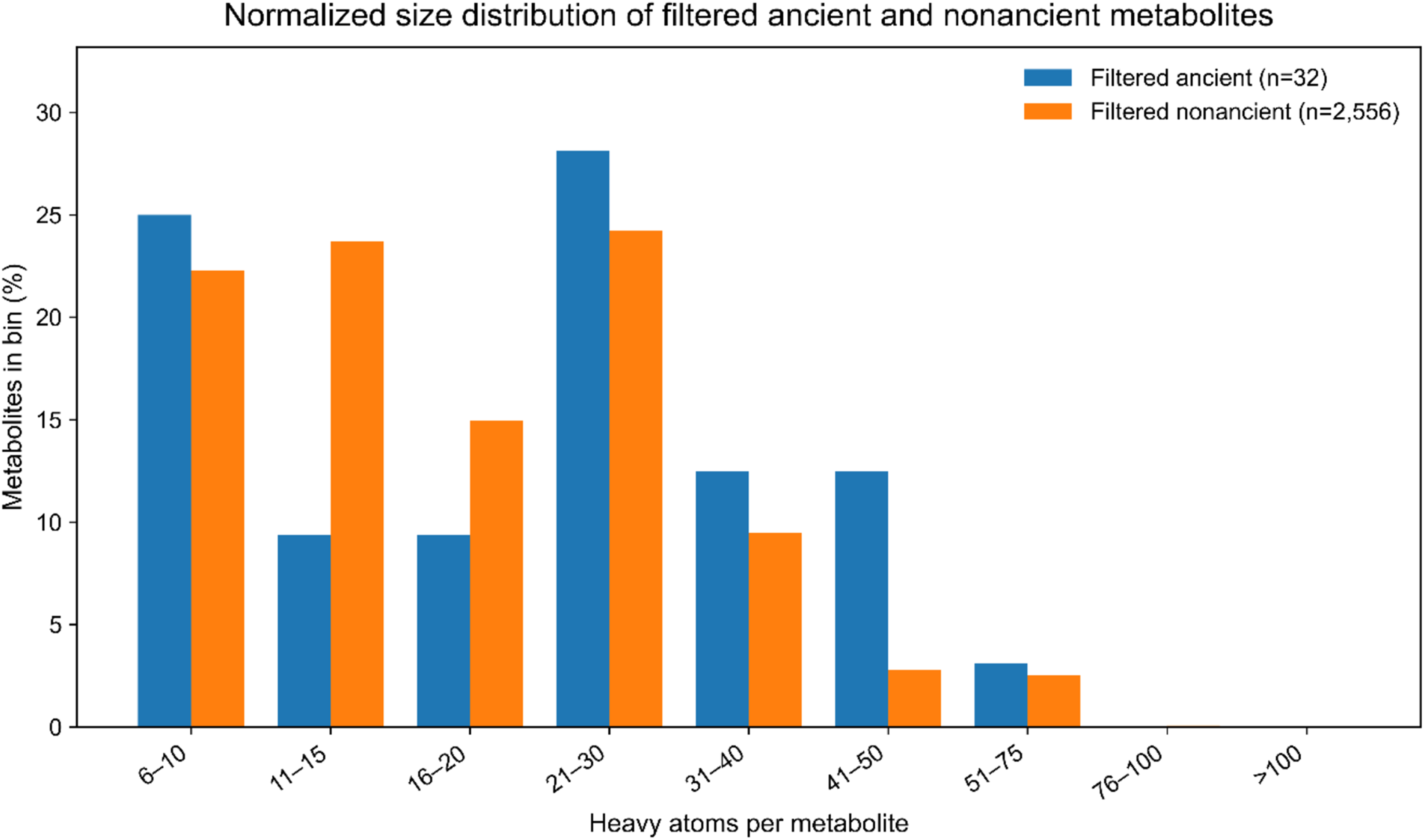
Normalized molecular-size distributions of ancient and filtered non-ancient metabolites. Distributions are shown as normalized heavy-atom-count frequencies for the stringent filtered sets after exclusion of BioLiP potential-artifact/dual-use ligands, elemental metals, and compounds containing fewer than six heavy atoms (n = 32 ancient; n = 2,556 non-ancient).

### Dense connectivity is not explained by close evolutionary relatedness

The median maximum sequence identities for filtered ancient versus filtered non-ancient edges were 35.2% versus 34.8% for one shared metabolite, 36.1% versus 35.5% for two, 36.6% versus 35.9% for three, 36.7% versus 36.1% for four, 37.0% versus 36.3% for five, and 37.0% versus 36.2% for exactly six shared metabolites. Thus, the ancient network already displays essentially the same sequence-diverse connectivity architecture observed after expansion of the full metabolite repertoire. Later chemical diversification appears to have expanded and densified an established network architecture rather than fundamentally changing its organization in protein-sequence space.

### Ancient metabolites are more strongly enriched for broad coverage of protein family space

Next, the exact BioLiP potential-artifact/dual-use ligand list was excluded, all elemental metal ions were removed from both metabolite classes, and metabolites containing fewer than six heavy atoms were excluded. The remaining 32 ancient metabolites occupied a mean of 185.38 ECOD F-groups per metabolite (median, 112), compared with 6.32 F-groups (median, 2) for the 2,540 filtered non-ancient metabolites mapped to ECOD. Ancient metabolites therefore exhibited a 29.35-fold enrichment in structural-family breadth. This enrichment was robust to bootstrap resampling (95% CI, 18.66–43.46) and highly significant by both Welch’s two-sample *t* test (16) (*t* = 4.63, df = 31.01, *P* = 6.22 × 10⁻⁵) and the Mann–Whitney *U* test (10) (*U* = 79,877, *P* = 7.56 × 10⁻²²). Thus, the extraordinary cross-family reuse of ancient metabolites cannot be attributed to elemental ions, small compounds, or the BioLiP potential-artifact/dual-use ligand class. Although this stringently filtered enrichment is lower than the initial unfiltered estimate, it provides a more conservative and defensible measure of the exceptional structural breadth of the ancient metabolite repertoire.

To test whether this exceptional breadth could arise by selecting the same number of non-ancient metabolites, we compared the complete 40-metabolite ancient set with 100,000 sets of 40 metabolites sampled without replacement from the artifact-filtered non-ancient pool. The observed ancient set had a mean breadth of 382.73 ECOD F-groups per metabolite (median, 142), whereas the random 40-metabolite sets had a mean of 4.52 (95% null interval, 2.30–9.83; median, 2). A heavy-atom-bin-matched null yielded a mean of 5.68 F-groups (95% null interval, 2.58–13.25; median, 2). None of the 100,000 unmatched or size-matched replicates reached the observed ancient mean (one-sided empirical P = 1.0 × 10⁻⁵ with the +1 correction). Moreover, 28 of the 40 ancient metabolites occupied at least 50 F-groups, compared with a mean of 0.64 in the size-matched null. The size of the average GCC is 2.3% of the allowed F-groups. Thus, the breadth of the ancient repertoire is not reproduced by arbitrary 40-member non-ancient sets, even after controlling for molecular-size composition.

### Combined ancient and non-ancient metabolite networks preserve global protein- family connectivity

As shown in Table 5, most F-groups represented in these networks are enzyme families, raising the possibility that the GCC arises trivially from extensive direct coupling among enzymes, i.e. the portion of the cellular network only associated with metabolism is globally connected. This analysis used the complete 40-metabolite ancient network, an artifact-filtered, metal-free non-ancient comparison set of 2,708 metabolites without the ≥6-heavy-atom cutoff, and the complete BioLiP2 metabolite set. These sets are therefore distinct from the stringently size-filtered ancient and non-ancient sets used for the 29.35-fold breadth comparison. Table 6 compares the unrestricted network with two constrained models: one allowing at most one direct enzyme– enzyme edge per enzyme and one prohibiting all direct enzyme–enzyme edges while retaining connections mediated through non-enzyme families.

**Table 5.** F-group composition by metabolite class.

| **Metabolite class** | **Total F-groups** | **Enzyme** | **Non-enzyme** | **Enzyme (%)** |
| --- | --- | --- | --- | --- |
| **40 ancient metabolites** | 6,798 | 6,596 | 202 | 97.03% |
| **Artifact-filtered, metal-free non-ancient (2,708; no size cutoff)** | 4,217 | 4,143 | 74 | 98.25% |
| **All BioLiP2 metabolites (42,106)** | 9,833 | 9,372 | 461 | 95.31% |

**Table 6.** Giant connected component comparison by enzymes and non-enzyme edge types.

| Metabolite class | Model | Total GCC | % of all F-groups | Enzymes in GCC | % enzyme F-groups |
| --- | --- | --- | --- | --- | --- |
| **Ancient** | 0: all edges allowed | 6,798 | 100.00% | 6,596 | 100.00% |
| **Ancient** | 1: $\leq 1$ direct E–E edge | 6,737 | 99.10% | 6,535 | 99.08% |
| **Ancient** | 2: no direct E–E edges | 6,691 | 98.43% | 6,489 | 98.38% |
| **Filtered non-ancient** | 0: all edges allowed | 4,187 | 99.29% | 4,113 | 99.28% |
| **Filtered non-ancient** | 1: $\leq 1$ direct E–E edge | 2,863 | 67.89% | 2,790 | 67.34% |
| **Filtered non-ancient** | 2: no direct E–E edges | 2,466 | 58.48% | 2,393 | 57.76% |
| **All metabolites** | 0: all edges allowed | 9,787 | 99.53% | 9,332 | 99.57% |
| **All metabolites** | 1: $\leq 1$ direct E–E edge | 9,474 | 96.35% | 9,019 | 96.23% |
| **All metabolites** | 2: no direct E–E edges | 9,237 | 93.94% | 8,782 | 93.70% |

As shown in Table 6, the ancient-metabolite network remained almost fully connected under both enzyme-edge restrictions, demonstrating that non-enzyme families can and likely extensively mediate indirect coupling among enzymes. The artifact-filtered, metal-free non-ancient set connected approximately two-thirds of its F-group network when each enzyme was allowed at most one direct enzyme–enzyme edge, whereas the complete BioLiP2 metabolite set remained almost fully connected. Thus, global connectivity is not driven solely by unrestricted direct enzyme–enzyme coupling; it can also arise through metabolite-mediated paths involving non-enzyme protein families. These results support a model in which later metabolite diversification expanded and elaborated a globally integrated architecture already generated by a small repertoire of ancient metabolites, irrespective of whether or not it is entirely associated with cellular metabolism.

## Discussion

The principal finding of this study is that ancient metabolites resolve an apparent paradox: they are simultaneously locally selective and globally connective. Many ancient metabolites preferentially occupy cognate ECOD F-groups or restricted structural families, demonstrating that repeated occurrence across the Protein Data Bank does not imply indiscriminate binding. After exclusion of the BioLiP potential-artifact/dual-use ligand class, elemental metals, and compounds containing fewer than six heavy atoms, ancient metabolites spanned protein-family space 29.35- fold more broadly per metabolite than filtered non-ancient metabolites. Collectively, the 32 filtered ancient metabolites connected all 3,135 associated ECOD F-groups into a single giant connected component. Thus, broad structural reach emerges not from a loss of specificity but from the repeated reuse of a limited metabolite vocabulary across diverse protein families. These findings suggest that selective molecular recognition and proteome-scale connectivity are complementary rather than opposing properties.

This distinction provides a structural framework for Entabolons, defined as groups of proteins whose functions become correlated through modulation by shared metabolites (3). At the molecular level, metabolites exhibit preferential binding to cognate folds and superfamilies. At the proteome level, however, extensive metabolite reuse generates overlapping binding repertoires that connect otherwise unrelated proteins and pathways. Within such a framework, fluctuations in metabolite abundance could produce coordinated changes in protein activity, conformation, localization, stability, or complex assembly, even among proteins that neither physically interact nor share a recent evolutionary origin. Fold specificity therefore provides the microscopic basis for a potential higher-order layer of metabolite-mediated functional organization.

The degree-preserving null model sharpens this interpretation. Near-complete giant-component occupancy was reproduced when the ECOD F-group network was randomized using the observed node-degree distribution. Across 250 configuration-model networks—50 replicates at each of five progressive metabolite-removal steps—the randomized GCC contained 100% of the F-groups in every replicate, matching the observed networks. The degree-cap analysis further showed that all 6,798 F-groups remained connected when each node was limited to only three edges. Consequently, the connectivity observed here does not require invocation of a specialized higher-order network architecture. Rather, it emerges naturally from the empirical degree distribution produced by the unusually broad reuse of ancient metabolites across structural families. The biological signal therefore lies not in exotic network wiring but in the extraordinary breadth with which a relatively small set of protein binding metabolites is deployed throughout protein-fold space.

Several important caveats should be considered. First, the present analysis treats metabolite binding largely as a binary event, whereas biological interactions occur across a continuum of affinities. Structural observations alone cannot distinguish among high-affinity cognate interactions, low-affinity regulatory interactions, and opportunistic binding events. Future analyses incorporating experimentally measured affinity distributions for representative systems such as NAD–Rossmann enzymes, thiamine diphosphate-dependent proteins, and GTPase families would provide a more nuanced understanding of the functional significance of metabolite reuse. Second, the study relies on PDB-derived structural observations, which inevitably reflect biases in protein selection, crystallization success, ligand availability, and community research priorities. Highly studied enzymes and cofactors are likely overrepresented, whereas some biologically relevant but transient interactions may be underrepresented. Although the scale of the analysis mitigates some of these limitations, the observed distributions should nevertheless be interpreted as a census of the current structural record rather than an unbiased survey of all biological interactions.

Third, the evolutionary implications of these findings remain provisional. Recurrent ligand-binding pockets may arise in part from the restricted set of geometries available when secondary-structure elements assemble into stable globular proteins (12). If such pockets emerged before extensive functional specialization, metabolites possessing favorable physicochemical properties could have been repeatedly recruited and subsequently retained as protein architectures diversified. The exceptional breadth of ancient-metabolite occupancy is consistent with this scenario, but the degree-preserving null model demonstrates that the giant connected component itself should not be interpreted as independent evidence for evolutionary templating. Rather, the evolutionary argument rests on the combination of metabolite selectivity, structural reuse, and fold-family breadth.

Although the observed differences in protein-family breadth and network connectivity are compelling, they may reflect several correlated properties of ancient metabolites, including their greater evolutionary age, central metabolic roles, higher cellular abundance, and repeated reuse across diverse biological processes. Thus, the data do not demonstrate that ancient metabolites possess an intrinsic mechanistic property that promotes connectivity. Rather, they show that metabolites retained since early evolution are highly enriched for molecules repeatedly recruited across protein-family space.

The analysis further distinguishes biologically meaningful breadth from trivial small-molecule promiscuity. Cross-fold scatter decreases with ligand size, and the highest-scatter ligands are predominantly very small crystallization additives. Similarly, ligand-balanced pocket- volume analyses demonstrate that generic additives preferentially occupy substantially smaller and more permissive cavities than metabolite-binding sites. These observations argue that much of the apparent promiscuity often attributed to metabolites can instead be explained by basic physicochemical constraints associated with molecular size and binding-site geometry. Importantly, after removal of artifact-prone ligands, elemental metals, and compounds containing fewer than six heavy atoms, the 32 ancient metabolites still occupied a mean of 185.38 ECOD F-groups per metabolite, compared with 6.32 for 2,540 mapped filtered non-ancient metabolites. The resulting 29.35-fold enrichment was robust to bootstrap resampling (95% CI, 18.66–43.46) and was not attributable to a significant difference in heavy-atom-count distributions.

Importantly, the present census is fundamentally correlational and does not establish functional coupling between proteins that share metabolites. Instead, it generates experimentally testable predictions for the Entabolon hypothesis (3). If overlapping metabolite-binding repertoires functionally couple proteins, then perturbation of a shared metabolite could induce coordinated responses among structurally unrelated proteins predicted to occupy the same metabolite-defined network. One approach to experimentally validate this idea would be to combine acute metabolite perturbation with proteome-wide thermal profiling, limited proteolysis, hydrogen–deuterium exchange mass spectrometry, or chemoproteomic methods to identify proteins whose conformational states change in concert. Orthogonal studies could measure enzymatic activity, protein-complex assembly, phase behavior, localization, or stability following controlled manipulation of metabolite abundance. Particularly informative systems would include canonical ancient metabolite–protein pairs such as NAD–Rossmann enzymes, thiamine diphosphate-dependent proteins, or GTPase networks, in which metabolite availability can be experimentally varied while responses are monitored across predicted network members. The strongest evidence for Entabolon behavior would be the observation that proteins sharing a metabolite exhibit correlated conformational or functional responses despite lacking direct physical interaction or close evolutionary relatedness. Causality should then be tested by editing metabolite- contact residues in two or more evolutionarily unrelated network members while preserving their expression and folding. Loss of the corresponding activity, localization, stability, or assembly response—and rescue by a conservative binding-competent allele—would connect metabolite binding to coordinated protein behavior. Time-resolved isotope tracing could further determine whether protein responses follow the shared-metabolite trajectory and precede changes in transcription. Finally, a minimal cell-free reconstitution containing several unrelated predicted binders would provide the most direct test: titration of one ancient metabolite should coordinately tune multiple proteins, whereas pocket mutants should sever individual response branches.

Experimentally determined coverage of bound metabolites in the PDB provides a conservative lower bound on metabolite-mediated connectivity. The present analysis was deliberately restricted to ECOD F-groups associated with experimentally observed protein–ligand interactions in BioLiP2 and the PDB. The stringently filtered combined ancient and non-ancient network spanned 5,388 of the 9,833 BioLiP-represented ECOD v295 F-groups, or 54.8% of this experimentally classified F-group universe; 5,367 of these families belonged to a single GCC. We did not infer additional metabolite–protein interactions using virtual ligand-screening approaches such as LIGMAP or related structure-based ligand-prediction methods (1, 4, 18, 19). Thus, the absence of an F-group from the present network should not be interpreted as evidence that its members cannot bind metabolites. Some unrepresented interactions may involve weak but functionally consequential binding, whereas others may be high-affinity interactions that have simply not yet been structurally or biochemically characterized. The emergence of a near-complete giant connected component within the experimentally supported portion of fold-family space therefore represents a conservative lower bound on the potential extent of metabolite-mediated protein-family connectivity. Extension of this framework to the complete PDB–ECOD fold space, integrating experimentally determined interactions with rigorously benchmarked computational ligand-binding predictions and prospective experimental validation, will determine whether metabolite-mediated connectivity extends across substantially more—and potentially most—of protein fold-family space.

The global connectivity of ancient metabolite networks may arise from the ability of individual metabolites to interact with many proteins. Approximately 500 geometrically distinct pocket types span the ligand-binding space of contemporary proteins, and these pockets may emerge largely from the physical constraints of compact, secondary-structure-containing folds rather than from independent optimization for every biochemical function (6, 20). Environmental cycling and phase-separated droplets could have generated and concentrated peptides approximately 40–100 residues long. Fluctuations toward homochirality would stabilize their secondary structures and compact folds, producing ligand-binding pockets and weak catalytic activities (21). Significantly, structurally generated peptide ensembles contain folds associated with ancient enzymes and pathways even without selection for specific functions, suggesting that physical constraints inherently bias early proteins toward metabolically useful structures (21).

A diverse population of these compact peptides could therefore have interacted promiscuously but selectively with ancient metabolites, creating globally connected protometabolic networks before the emergence of highly specific enzymes. Weak proto-ligases may have further expanded this population by joining shorter peptides, establishing positive feedback among peptide production, folding, ligand binding, and catalysis. Natural selection could subsequently refine this physically generated ensemble into modern enzymes and regulated pathways. Thus, ancient metabolites may have acted not as indiscriminate binders but as selective, repeatedly used molecular connectors organizing protein-fold space into an almost completely connected network—the central premise of the Entabolon hypothesis.

Taken together, these results support a simple but potentially powerful principle: ancient metabolites are not indiscriminate binders but selective, highly reused molecular connectors. Their empirical binding distributions are sufficient to organize protein-fold space into an almost completely connected metabolite-mediated network. Whether this structural connectivity translates into dynamic, metabolite-driven coordination of protein behavior remains an open question, but the present work provides both the quantitative framework and the experimentally testable predictions needed to evaluate the Entabolon hypothesis directly.

## Acknowledgments

This research was supported in part by a gift from the Parker H. Petit AI-Driven Drug Discovery Initiative and a grant GM-18039 from the National Institute of General Medical Sciences of the National Institutes of Health. The authors declare no competing interests. We thank Jessica Forness for reviewing this manuscript and Bartosz Ilkowski for his computational support.

## Data and code availability

All primary data derive from public resources (BioLiP2, ECOD v295, the PDB Chemical Component Dictionary, HMDB, and ChEBI). Derived datasets and analysis code will be deposited in a public repository before publication and are available from the corresponding author upon reasonable request.

## Methods

### ECOD and BioLIP2 analysis

The full 2026 BioLiP2 annotation set, comprising 989,058 binding sites (12, 13), was joined to ECOD v295 domains (5) by residue-range attribution. Each binding site was assigned to the ECOD H-group of the domain containing the majority of its binding residues; 929,546 of 989,058 sites (94.0%) received an H-group assignment. H-groups were used for site-level specificity and scatter analyses because they capture homologous superfamilies while avoiding fragmentation among closely related families. Specificity was defined as the fraction of assigned sites belonging to the modal H-group. Cross-fold scatter was defined as the effective number of occupied H-groups, calculated as exp(Shannon entropy) (16), and was rarefied to 50 sites by multivariate-hypergeometric subsampling using 30 replicates. Ligand size was defined as the nonhydrogen atom count obtained from the PDB Chemical Component Dictionary (17). Ligands were classified as ancient-reference metabolites, other biological ligands, or small crystallization additives. The 40 ancient metabolites represented in the HMDB/ChEBI–BioLiP–ECOD intersection are listed in Table S1 and were used for the complete ancient-network analyses. For stringent ancient–non-ancient comparisons, BioLiP potential-artifact/dual-use ligands, elemental metals, and compounds containing fewer than six heavy atoms were excluded, leaving 32 ancient metabolites. Specificity and scatter analyses were restricted to ligands with at least 20 assigned sites; generic peptide, RNA, and DNA ligands were excluded. Associations were evaluated using Spearman’s ρ (9), and between-class comparisons were performed using two-sided Mann– Whitney *U* tests (10).

### Identification of metabolites and fold-family breadth

BioLiP2 protein–ligand annotations were mapped independently to ECOD v295 F-groups at the PDB-chain level (16–18). F-groups were used for breadth and network analyses because the questions addressed here concern the number and connectivity of distinct structural families rather than site-level superfamily preference. The metabolite background comprised PDB Chemical Component Dictionary entries mapping to HMDB structures or ChEBI entities classified under the metabolite-role ontology (19, 20). Components were matched to HMDB by InChIKey, and ChEBI metabolite annotations were obtained from the ChEBI relation and name tables. Forty ancient metabolites were represented in the HMDB/ChEBI–BioLiP–ECOD intersection and were used for the complete ancient-network analyses. Fold-family breadth was defined as the number of distinct ECOD F-groups represented among BioLiP2 PDB chains binding a metabolite.

For the stringent ancient–non-ancient breadth comparison, the BioLiP potential- artifact/dual-use ligand list was applied, elemental metal ions were removed from both classes, and metabolites containing fewer than six heavy atoms were excluded. This filtering retained 32 ancient metabolites and 2,556 non-ancient metabolites, of which 2,540 mapped to ECOD F-groups. The enrichment factor was calculated as the mean breadth of the 32 ancient metabolites—185.38 F-groups per metabolite—divided by the mean breadth of the 2,540 mapped filtered non-ancient metabolites—6.32 F-groups per metabolite—yielding a 29.35-fold enrichment. Confidence intervals were obtained by independently resampling metabolites with replacement within each class for 10,000 bootstrap replicates.

For the F-group networks, nodes represented ECOD F-groups, and edges joined two families when they shared the specified minimum number of metabolites from the analyzed class. The complete network defined by the 40 ancient metabolites contained 6,798 F-groups, whereas the stringently filtered ancient, stringently filtered non-ancient, and combined stringently filtered networks contained 3,135, 4,087, and 5,388 F-groups, respectively. Site-level H-group assignments used the independent ECOD v295 residue-range join described above. Network coverage was contextualized against the 9,833 BioLiP-represented ECOD v295 F-groups.

### Degree-preserving network null model

The ancient-metabolite ECOD F-group network was reconstructed from BioLiP2 using the 40 ancient-metabolite CCD codes represented in the fold-breadth analysis. Duplicate PDB-chain– ligand associations were collapsed, BioLiP2 chains were mapped to ECOD v295 F-groups, and each metabolite was assigned the set of distinct F-groups represented among its binding chains. The one-mode family network contained 6,798 F-group nodes and 10,011,118 unique edges, with an edge joining two F-groups when they bound at least one common ancient metabolite.

Two degree-preserving null analyses were performed at different network levels. For the protein-chain bipartite graph, null networks were generated by swapping metabolite endpoints between pairs of chain–metabolite edges only when the resulting edges were absent, thereby preserving exactly the degree of every protein chain and metabolite in a simple bipartite graph. A total of 412 chain-level randomizations were analyzed. For the ECOD F-group one-mode network, null networks were generated directly from the observed family-degree distribution using a degree-preserving configuration model. Node stubs were randomly paired, preserving each F-group degree exactly; self-loops and parallel edges were permitted in the null multigraph. Fifty independent family-level randomizations were performed at each of five metabolite-removal stages—the complete 40-metabolite set, removal of five elemental metals, additional removal of ATP/ADP, additional removal of NAD/NADP, and additional removal of GTP/GDP—for a total of 250 family-level randomized networks. GCC fractions were calculated among nodes represented after each removal stage, and observed values were compared with the corresponding empirical null distributions using two-sided empirical P values. All completed randomized networks were analyzed without further selection.

## Supplementary Information

**Table S1.** Demonstration that the promiscuous small-ligand tail is heterogeneous^a^.

| Ligand (CCD) | # of heavy atoms | Character | Functional role / binds preferentially to |
| --- | --- | --- | --- |
| Glycerol (GOL) | 6 | artifact (precipitant) | none |
| Ethylene glycol (EDO) | 4 | artifact (precipitant) | none |
| PGE (PGE) | 10 | artifact (precipitant) | none |
| PEG fragment (PEG) | 7 | artifact (precipitant) | none |
| 2-methyl-2,4-pentanediol (MPD) | 8 | artifact (precipitant) | none |
| Dimethyl sulfide (DMS) | 3 | artifact / nonspecific | none |
| Dimethyl sulfoxide (DMSO) | 4 | artifact / solvent | none |
| Urea (URE) | 4 | nonspecific / denaturant | none |
| Sulfate (SO4) | 5 | dual-function | allosteric anion effector (Mode 4); phosphate-mimic at anion pockets |
| Phosphate (PO4) | 5 | dual-function | phosphate-binding pockets; functional more often than not |
| Pyrophosphate (PPi) | 9 | dual-function | phosphate / anion pockets; functional |
| Imidazole (IMD) | 5 | broad-family binder | His / metal (His-tag) sites |
| Citrate (CIT) | 13 | dual-function | buffer / chelator + allosteric effector (e.g. PFK) |
| Acetate (ACT) | 4 | broad-family / metabolite | oxyanion holes, acyl pockets |
| Formate (FMT) | 3 | broad-family / metabolite | anion sites |
| Pyruvate (PYR) | 6 | small metabolite | various |
| Succinate (SIN) | 8 | small metabolite | dicarboxylate sites |
| Fumarate (FUM) | 8 | small metabolite | dicarboxylate sites |
| Glycine (GLY) | 5 | small metabolite | various |

<sup>a</sup> True precipitants (top) organize no defined protein module. A dual-function group—including sulfate, phosphate, pyrophosphate, and citrate—combines crystallization prevalence with genuine functional binding; sulfate also recurs as an allosteric anion effector in the companion distal-effector analysis. Broad-family binders and genuine small metabolites complete the high-scatter tail. Thus, cross-fold scatter must be mechanistically decomposed rather than equated with artifact.

**Table S2.** Ancient-metabolite set and fold and functional properties^a^.

| Metabolite | Cognate / favored fold | Specificity | Scatter | # of heavy atoms | Sub-class |
| --- | --- | --- | --- | --- | --- |
| **NAD** | Rossmann (NAD-binding) | 0.76 | 2.6 | 44 | redox cofactor |
| **NADP** | Rossmann (dehydrogenase) | 0.63 | 3.5 | 48 | redox cofactor |
| **FAD** | FAD-binding (Rossmann) | 0.50 | 5.7 | 53 | redox cofactor |
| **FMN** | flavodoxin / TIM-barrel | 0.29 | 8.0 | 31 | redox cofactor |
| **Riboflavin** | flavin-binding | 0.24 | 8.6 | 27 | redox cofactor |
| **Heme b** | globin / cytochrome / P450 | 0.24 | 10.5 | 43 | redox cofactor |
| **Lipoate** | lipoyl / acyltransferase | n/a | n/a | 12 | redox cofactor |
| **Glutathione** | thioredoxin / GST | 0.72 | 3.0 | 20 | redox cofactor |
| **Ascorbate** | various | n/a | n/a | 12 | redox cofactor |
| **ATP** | P-loop NTPase / kinase | 0.54 | 6.9 | 31 | nucleotide |
| **ADP** | P-loop NTPase / kinase | 0.49 | 5.9 | 27 | nucleotide |
| **AMP** | mononucleotide-binding | 0.10 | 21.3 | 23 | nucleotide |
| **GTP** | P-loop / TRAFAC GTPase | 0.65 | 3.6 | 32 | nucleotide |
| **GDP** | P-loop NTPase | 0.62 | 2.6 | 28 | nucleotide |
| **GMP** | mononucleotide-binding | 0.17 | 12.9 | 24 | nucleotide |
| **cAMP** | cyclic-nucleotide-binding | n/a | n/a | 22 | nucleotide |
| **CoA** | multiple (broad) | 0.18 | 12.2 | 48 | carrier |
| **Acetyl-CoA** | acyltransferase (broad) | 0.41 | 7.5 | 51 | carrier |
| **SAM** | Rossmann (methyltransferase) | 0.49 | 5.2 | 27 | carrier |
| **SAH** | Rossmann (methyltransferase) | 0.77 | 2.3 | 26 | carrier |
| **Biotin** | biotin/lipoyl-attachment | 0.63 | 3.5 | 16 | carrier |
| **Thiamine diphosphate** | thiamine-diphosphate fold | 0.98 | 1.1 | 26 | carrier |
| **PLP** | PLP-dependent transferase | 0.64 | 3.3 | 16 | carrier |
| **Pyruvate** | various | 0.54 | 5.9 | 6 | intermediate |
| **2-oxoglutarate** | DSBH dioxygenase | 0.64 | 3.9 | 10 | intermediate |
| **Citrate** | metal/anion sites (broad) | 0.36 | N<50 | 13 | intermediate |
| **Succinate** | dicarboxylate sites | N<20 | N<20 | 8 | intermediate |
| **Fumarate** | dicarboxylate sites | 0.39 | 6.9 | 8 | intermediate |
| **Phosphoenolpyruvate** | PEP-utilizing (broad) | 0.82 | 2.3 | 10 | intermediate |
| **Fructose-1,6-bisphosphate** | sugar-phosphate / allosteric | 0.74 | 2.7 | 20 | intermediate |
| **Glycine** | various (broad) | 0.23 | 15.1 | 5 | amino acid |
| **Alanine** | various (broad) | 0.20 | 15.6 | 6 | amino acid |
| **Aspartate** | various (broad) | 0.34 | 8.9 | 9 | amino acid |
| **Glutamate** | various (broad) | 0.45 | 6.8 | 10 | amino acid |
| **Serine** | various (broad) | 0.20 | 13.9 | 7 | amino acid |
| **Mg<sup>2+</sup>** | metal-dependent enzymes | 0.09 | 25.8 | 1 | metal |
| **Zn<sup>2+</sup>** | metalloenzymes / Zn-finger | 0.10 | 24.5 | 1 | metal |
| **Mn<sup>2+</sup>** | metal-dependent enzymes | 0.12 | 19.3 | 1 | metal |
| **Ca<sup>2+</sup>** | Ca-binding / signaling | 0.10 | 26.8 | 1 | metal |
| **Fe (ion)** | metalloenzymes | 0.43 | 9.3 | 1 | metal |

<sup>a</sup> Specificity is the fraction of sites in the modal ECOD H-group; scatter is the effective number of H-groups, rarefied to 50 sites over 30 replicates. Entries marked N<20 or N<50 contain too few assigned sites for a stable estimate; n/a indicates that the CCD entry was not resolved in this census. Table S3 lists the exact 40 ancient metabolites represented in the final HMDB/ChEBI–BioLiP–ECOD intersection used for F-group breadth and network analyses.

**Table S3.** Ancient metabolites represented in the fold-breadth analysis^a^.

| Metabolite | PDB CCD code | ECOD fold families | BioLiP PDB chains |
| --- | --- | --- | --- |
| 2-oxoglutarate | AKG | 107 | 856 |
| ADP | ADP | 919 | 11,979 |
| AMP | AMP | 440 | 1,871 |
| ATP | ATP | 921 | 8,598 |
| Acetyl-CoA | ACO | 120 | 725 |
| Alanine | ALA | 101 | 261 |
| Ascorbate | ASC | 33 | 64 |
| Aspartate | ASP | 81 | 433 |
| Biotin | BTN | 26 | 380 |
| Ca<sup>2+</sup> | CA | 2,404 | 27,383 |
| Citrate | CIT | 10 | 21 |
| CoA | COA | 225 | 1,794 |
| FAD | FAD | 254 | 6,751 |
| FMN | FMN | 175 | 3,673 |
| Fe (ion) | FE | 316 | 6,029 |
| Fructose-1,6-bisphosphate | FBP | 25 | 669 |
| Fumarate | FUM | 45 | 134 |
| GDP | GDP | 260 | 8,186 |
| GMP | GMP | 31 | 78 |
| GTP | GTP | 284 | 8,434 |
| Glutamate | GLU | 164 | 1,125 |
| Glutathione | GSH | 94 | 1,043 |
| Glycine | GLY | 204 | 833 |
| Heme b | HEM | 207 | 14,069 |
| Lipoate | LPA | 5 | 15 |
| Mg<sup>2+</sup> | MG | 2,872 | 40,217 |
| Mn<sup>2+</sup> | MN | 865 | 9,388 |
| NAD | NAD | 291 | 6,023 |
| NADP | NAP | 213 | 4,321 |
| PLP | PLP | 81 | 2,933 |
| Phosphoenolpyruvate | PEP | 32 | 298 |
| Pyruvate | PYR | 91 | 529 |
| Riboflavin | RBF | 27 | 124 |
| SAH | SAH | 242 | 2,721 |
| SAM | SAM | 234 | 1,284 |
| Serine | SER | 117 | 257 |
| Succinate | SIN | 10 | 9 |
| Thiamine diphosphate | TPP | 40 | 959 |
| Zn<sup>2+</sup> | ZN | 2,696 | 52,656 |
| cAMP | CMP | 47 | 393 |

<sup>a</sup> Forty ancient metabolites were represented in the final HMDB/ChEBI–BioLiP–ECOD intersection used for the complete ancient-network analysis. For the stringent breadth comparison, BioLiP potential-artifact/dual-use ligands, elemental metal ions, and compounds containing fewer than six heavy atoms were excluded. This retained 32 ancient metabolites and 2,556 non-ancient metabolites, of which 2,540 mapped to ECOD F-groups. Fold-family breadth was the number of distinct ECOD v295 F-groups represented among BioLiP2 PDB chains binding a metabolite. Mean breadth was 185.38 F-groups for the 32 ancient metabolites and 6.32 for the 2,540 mapped filtered non-ancient metabolites, yielding 29.35-fold enrichment (bootstrap 95% CI, 18.66–43.46; 10,000 within-class bootstrap replicates). Network nodes represented ECOD F-
groups, and edges joined families sharing the specified minimum number of metabolites. Network coverage was contextualized against the 9,833 BioLiP-represented ECOD v295 F-groups.

